# Sonogenetics for noninvasive and cellular-level neuromodulation in rodent brain

**DOI:** 10.1101/2020.01.28.919910

**Authors:** Yaoheng Yang, Christopher Pham Pacia, Dezhuang Ye, Lifei Zhu, Hongchae Baek, Yimei Yue, Jinyun Yuan, Mark J. Miller, Jianmin Cui, Joseph P. Culver, Michael R. Bruchas, Hong Chen

**Author notes:** Address correspondence to Hong Chen, Ph.D. Department of Biomedical Engineering and Radiation Oncology, Washington University in St. Louis. 4511 Forest Park Ave. St. Louis, MO, 63108, USA.

## Abstract

Sonogenetics, which uses ultrasound to noninvasively control cells genetically modified with ultrasound-sensitive ion channels, can be a powerful tool for investigating intact brain circuits. Here we show that TRPV1 is an ultrasound-sensitive ion channel that can modify the activity of TRPV1-expressing cells *in vitro* when exposing to focused ultrasound. We also show that focused ultrasound exposure at the mouse brain *in vivo* can selectively activate neurons that are genetically modified to express TRPV1. We demonstrate that precise manipulation of neural activity via TRPV1-based sonogenetics can be achieved by spatiotemporal control of ultrasound-induced heating. The focused ultrasound exposure is safe based on our inspection of neuronal integrity, apoptosis, and inflammation markers. This sonogenetic tool enables noninvasive, cell-type specific, spatiotemporally controlled modulation of mammalian brain activity.

## Introduction

Technologies for noninvasive modulation of mammalian brain activity with spatiotemporal precision and cell-type specificity have long been desired for experimental investigation of intact neural systems. Compared with electric, magnetic, optical, and chemical stimuli that have been used by existing neuromodulation technologies, focused ultrasound (FUS) offers unique advantages as it can noninvasively deliver ultrasound energy through the intact human scalp and skull deep into the brain and focus at cortical areas^1, 2^ as well as deep brain areas^3^ with high spatial selectivity. Existing FUS neuromodulation techniques use ultrasound pulses with low intensity to produce mechanical effects for neural and behavioral modulation with a negligible temperature increase^4^. Numerous studies have confirmed that FUS can noninvasively modulate neural activity and brain function in animal models^5, 6^ and humans^1, 3^. However, the broad application of FUS neuromodulation is challenged by the wide variation in neural cell types in the sonicated brain tissue, resulting in neuromodulation with relatively low reliability and replicability^7, 8^. Advances in genetics-based tools enable manipulation of a specific type of neurons embedded within densely-wired brain circuits for assessing the causal role that different groups of neurons have in controlling circuit activity and behavior outcomes. Among genetics-based tools, optogenetics has been transforming neuroscience research by allowing targeted control of precisely defined events in the brain via light stimulation; however, it requires invasive surgery to permanently implant devices for delivering light deep into the brain.

Similar to optogenetics, sonogenetics intends to use ultrasound to precisely control the activity of neurons that have been genetically modified to express ultrasound-sensitive ion channels (*i.e.*, sonogenetics actuators). The major challenge in the development of sonogenetics is to find suitable actuators. Sonogenetics was initially proposed by Ibsen *et al*^9^ where ultrasound selectively activated the cells expressing a mechanosensitive ion channel (TRP-4) in *Caenorhabditis elegans*. However, ultrasound alone did not affect the behavior of the *worm*. Behavioral changes were observed only when the body of the worm was surrounded by microbubbles added to the agar plate where the worm was placed. The requirement for adding microbubbles to activate the mechanosensitive signaling prevents the translation of this method to studies in the mammalian brain. A recent study using *Caenorhabditis elegans* found that the mechanosensitive ion channel MEC-4 was important for ultrasound neuromodulation in the absence of microbubbles^10^. But, its application in mammalian brains is limited because high-frequency ultrasound (10 MHz) used in that study cannot efficiently penetrate through the skull. Other mechanosensitive ion channels, for example, TREK-1^11^, MscL^12^, Piezo1^13, 14^, and TRPA1^15^ have been proposed as potential sonogenetic tools based on *in vitro* cell culture studies without *in vivo* validation. A most recent study showed that an engineered auditory-sensing protein (prestin) caused neurons in the mouse brain to sense ultrasound stimulation^16^; however, this protein, which is not an ion channel, lacks the ability to control neuron activity in a temporally precise fashion. To our knowledge, there is so far no study that has identified ultrasound-sensitive ion channels for spatiotemporally precise modulation of neural activity in mammalian brains.

Ultrasound propagation in tissue generates not only mechanical effects but also thermal effects owing to viscous dissipation as molecules move back and forth in the ultrasound field. Here, we present a sonogenetic tool (Fig. 1) for noninvasive, spatiotemporally controlled manipulation of cells genetically modified to express the thermosensitive ion channel – transient receptor potential vanilloid 1 (TRPV1). The activation temperature of TRPV1 is around 42 °C^17^, which is only a few degrees higher than the body temperature to permit quick and safe stimulation while allowing the channels to be closed at physiological temperature and minimizing other potential thermal effects on the neural circuits^18–20^. This technique opens new avenues for noninvasive, cell-type specific, spatiotemporally precise control of mammalian neural activity.

**Fig. 1.**
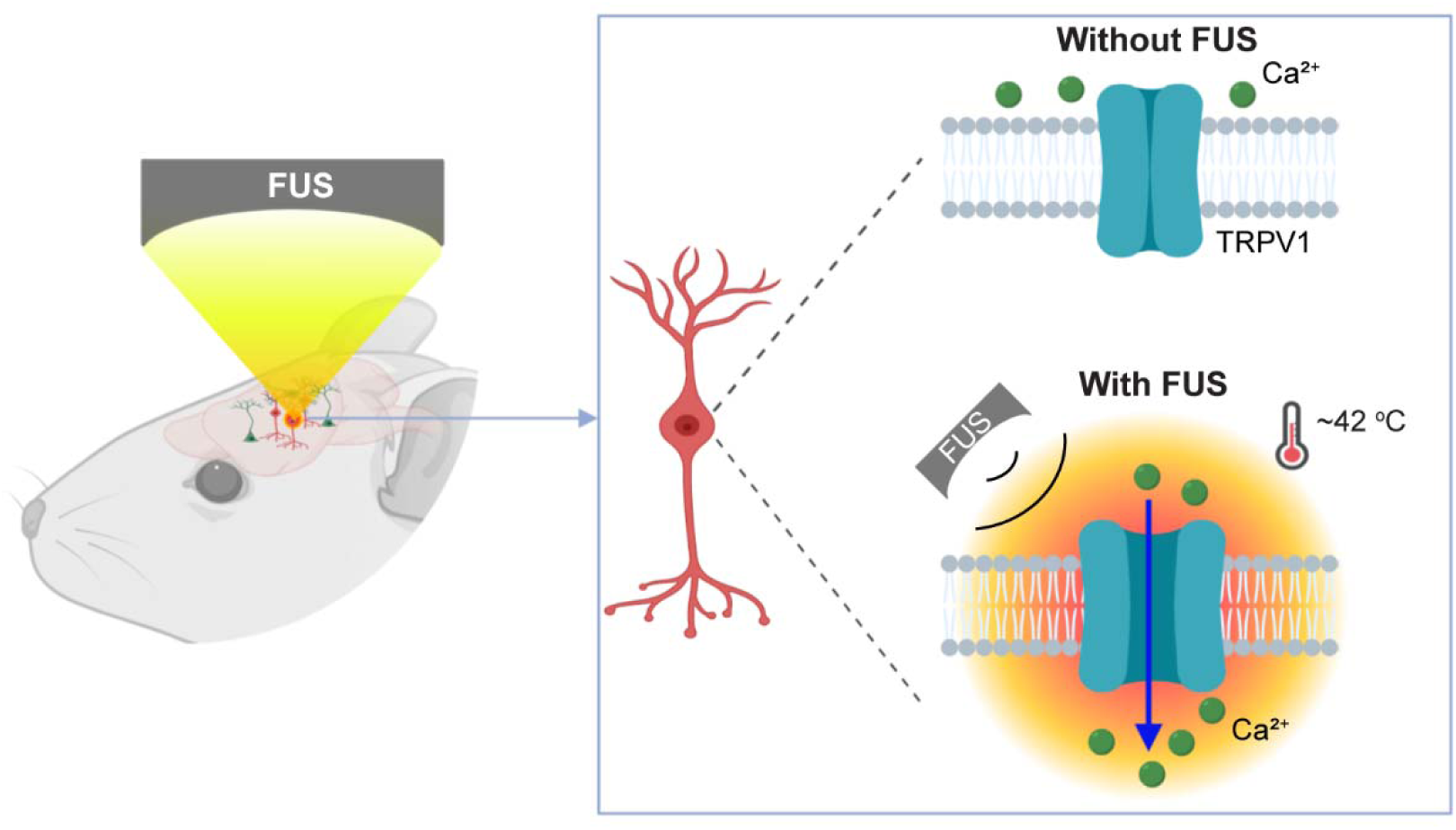
Schematic illustration of TRPV1-mediated sonogenetics. FUS directly and selectively activates neurons expressing the thermosensitive ion channel TRPV1 by spatiotemporally controlling the heating of targeted brain tissue to ~42 °C.

## Results

### TRPV1 is a sonogenetic actuator

Our first experiment was performed to determine whether TRPV1 is a sonogenetic actuator by evaluating whether ultrasound could selectively control intracellular Ca^2+^ influx in TRPV1-expressing HEK293T cells. We developed an experimental setup that allows simultaneous fluorescence imaging and FUS stimulation of HEK293T cells. This setup used a ring-shape FUS transducer that was placed underneath the cell culture plate, allowing fluorescence imaging through the center opening of the ring (Fig. S1). Temperature rise associated with the ultrasound exposure was measured using a fiber-optic thermometer placed ~1 mm away from the center point of the microscope’s field of view (FOV). The maximum temperature induced by FUS as recorded by the thermometer was 42.0 ± 0.2 ^°^C. The functionality of TRPV1 was confirmed by the observation of Ca^2+^ influx in response to capsaicin, a TRPV1 agonist (Fig. S2).

A high-amplitude Ca^2+^ influx was detected in the FUS-stimulated TRPV1-expressing cells (TRPV1^+^) using fluorescence imaging (Fig. 2a, 2c). The maximum increase in the fluorescence intensity(ΔF/F) was 0.46 on average (Fig. 2e). The TRPV1 antagonist Capsazepine^21, 22^ dramatically reduced Ca^2+^ influx in response to FUS stimulation in TRPV1^+^ cells, suggesting that the observed Ca^2+^ influx was due to TRPV1 activation (Fig. S3). To determine whether FUS stimulation affected cells not expressing TRPV1, we stimulated cells that did not express TRPV1 (TRPV1^-^) and did not evoke significant Ca^2+^ influx (Fig. 2c, 2d). In both TRPV1^-^ and TRPV1^+^ cells, there were no significant changes in intracellular Ca^2+^ concentration without FUS stimulation. FUS stimulation had no detectable effects on cell viability (Fig. S4). These findings suggest that TRPV1 can serve as a sonogenetic actuator.

**Fig. 2.**
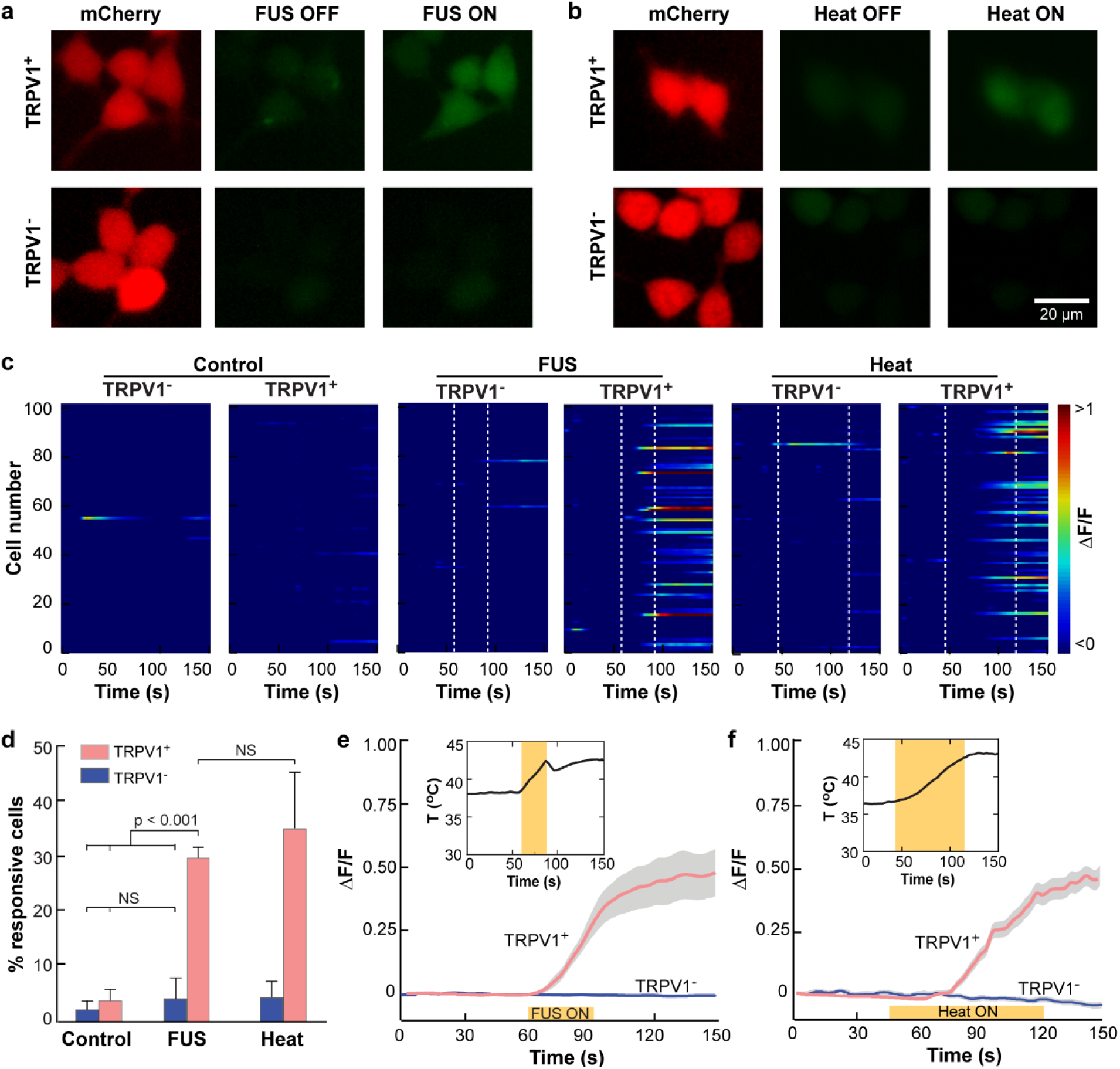
TRPV1 enables FUS activation of HEK293T cells *in vitro*. (**a, b**) Fluorescence images of HEK293T cells. Red: HEK293T cells expressing mCherry with TRPV1 ion channel (TRPV1^+^, top row) or without TRPV1 ion channel (TRPV1^-^, bottom row). Green: Intracellular Ca^2+^ before and after (**a**) FUS stimulation and (**b**) water-bath heating. (**c**) Heatmap of normalized Ca^2+^ fluorescence intensity change (ΔF/F) of 100 randomly selected TRPV1^+^ and TRPV1^-^ cells from 3 independent trials in the control group without any stimulation (left panel), FUS group (middle panel), and heating group (right panel), respectively. (**d**) Percentage of TRPV1^+^ and TRPV1^-^ cells within the field of view of the fluorescence microscope that responded to FUS stimulation and water-bath heating, respectively. Cells in the control group were not stimulated. The error bar indicates the standard error of the mean (SEM). Normalized fluorescence intensity change (ΔF/F) of TRPV1^+^ and TRPV1^-^ induced by (**e**) FUS stimulation and (**f**) water bath heating. The solid lines indicate the mean and shaded gray represents SEM. The insert shows the corresponding temperature curve.

To test our hypothesis that FUS activates the TRPV1^+^ cells through FUS thermal-dominated effects, we performed an experiment to heat the TRPV1^+^ cells by water bath heating. We controlled the heating duration to increase the temperature to 42.2 ± 0.5 ^°^C, which was approximately the same level as FUS-induced heating. We found that water-bath heating on TRPV1^+^ cells had a similar success rate and mean fluorescence intensity as FUS stimulation (Fig. 2). While we cannot rule out the contribution by the ultrasound mechanical effect, these findings suggest that the activation of the TRPV1 ion channel is dominated by the ultrasound thermal effect.

### Sonogenetics in the mouse brain

After establishing TRPV1 as a sonogenetic actuator through the *in vitro* experiment, we then tested whether FUS could directly activate TRPV1^+^ neurons in the mouse brain *in vivo.* We used a two-photon microscope (2PM) combined with genetically encoded calcium indicator (GCaMP6f) for *in vivo* neural activity recording in the mouse cortex. Calcium imaging allows direct observation of brain activity and overlaying functional activity with genetic and anatomical identity^23^. We intentionally avoided using electrophysiological recording because electrodes inserted in the brain interfere with ultrasound wave propagation, and ultrasound wave-induced mechanical vibration generates artifacts in the electrical recording.

The TRPV1 transgene was placed under the excitatory neuronal promotor calmodulin α-subunit and before mCherry by posttranscriptional cleavage linker p2A (CaMKII-TRPV1-p2A-mCherry)^20^. The transgene was packed into a lentiviral vector. Two groups of genetically engineered GCaMP6f mice were used in this experiment. Group one (n=4) was injected with the above lentiviral vector at the somatosensory cortex, and group two (n=4) was injected with a lentiviral vector encoding mCherry without TRPV1 at the same brain location (CaMKII-mCherry, Fig. 3a). After approximately 4 weeks for transgene expression, a cranial window was created on top of the virus injection site and a glass window was added for 2PM imaging of the neural activity by the genetically encoded calcium sensor GCaMP6f. A ring-shape FUS transducer was coupled to the microscope objective through a custom-designed adaptor for simultaneous FUS stimulation and 2PM imaging (Fig. 3a, 3b). The full-width-half-maximum dimension of the FUS beam was 0.66 mm in the horizontal plane (XY plane, Fig. 3a), which enables precise spatial control of the delivered FUS energy within the FOV of the 2PM microscope.

**Fig 3.**
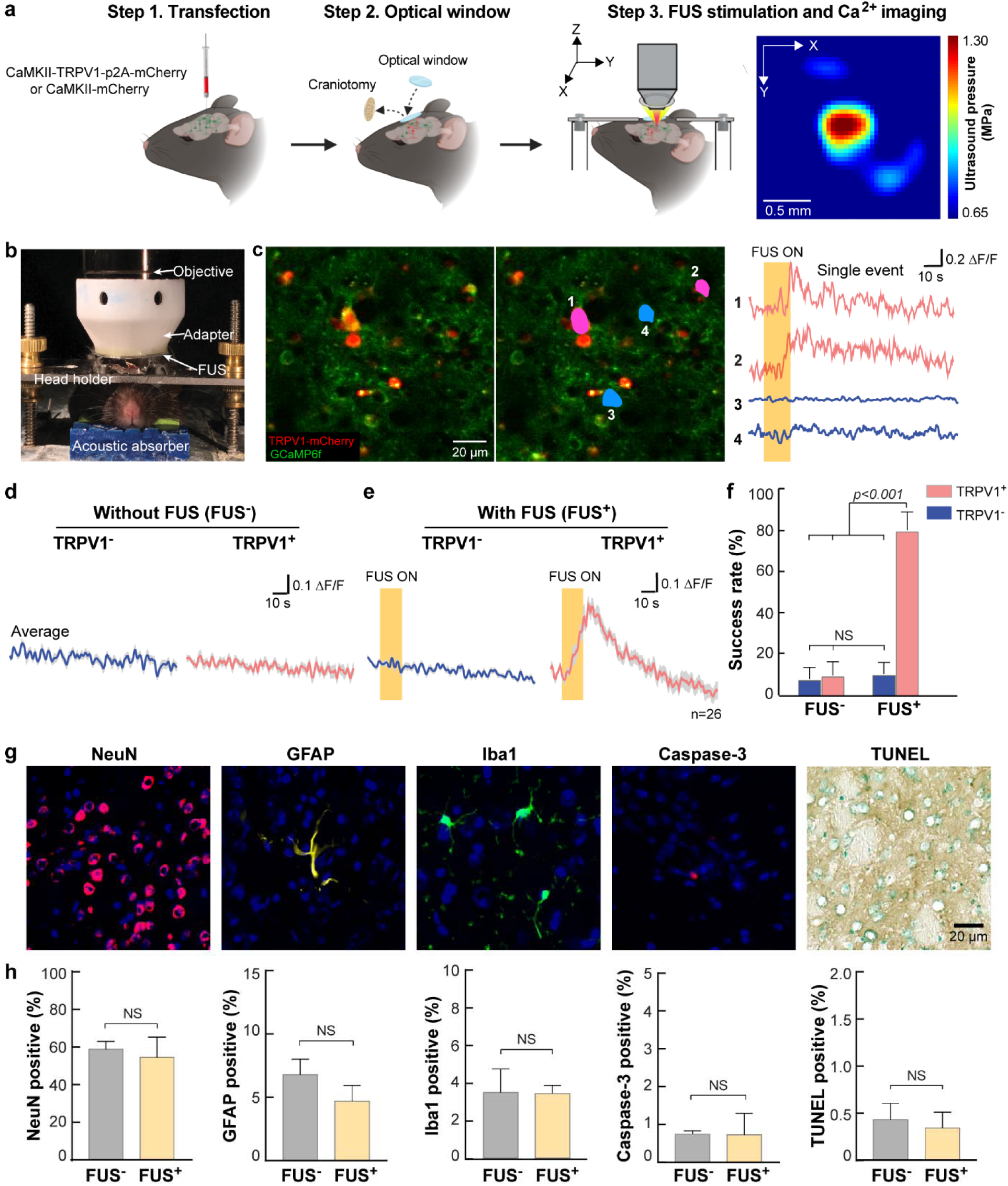
Sonogenetics in mouse brain *in vivo.* **(a)** Schematic illustration of the experiment workflow. The FUS pressure distribution in the 2PM imaging plane as measured by a hydrophone is shown on the right. (**b**) A photo of the 2PM setup which couples a ring-shaped FUS transducer with the microscope objective using a customized adapter. The mouse head was fixed by a holder to minimize motion artifact in 2PM imaging. An acoustic absorber was placed underneath the mouse head to decrease the reflection of the ultrasound pulses from the bottom of the mouse head. (**c**) A representative 2PM image of the mouse cortex *in vivo* is shown on the left. The image on the right highlights two neurons (1 and 2 in pink) expressing both TRPV1-mCherry and GCaMP6f and two neurons (3 and 4 in blue) expressing GCaMP6f only. The Ca^2+^ fluorescence intensity changes over time of the 4 neurons are shown next to the 2PM image. A total of 13 neurons were selected for mice with and without overexpression of TRPV1 and each neuron was stimulated by two repeated FUS stimulations. (**d**) Average of the Ca^2+^ fluorescence intensity curves for mice with and without overexpressing of TRPV1 before FUS exposure, which served as the control for without FUS stimulation (FUS^-^). (**e**) Average of the Ca^2+^ fluorescence intensity curves for mice with and without overexpression of TRPV1 with FUS stimulation (FUS^+^) Solid lines are the mean of the 26 stimulations and the shaded area indicates the SEM. **(f)** Comparison of the success rate of these selected neurons with and without FUS stimulation. The error bar represents SEM. **(g)** Evaluation of neuronal integrity, inflammation, and apoptosis after FUS exposure in the FUS-targeted location using immunohistochemical staining of the neurons (NeuN), astrocytes (GFAP), microglia (iba1), Caspase-3, and TUNEL. In the fluorescence images, blue indicates the DAPI stained nuclei and the other pseudo colors indicate different cell types. (**h**) Percentages of positive-stained cells to DAPI cells comparing without and with FUS exposure within the FUS targeted location.

We found a total of 13 neurons with co-expression of TRPV1 and GCaMP6f from mice in group one. Two repeated FUS stimulations were delivered when these neurons were found in the FOV. We observed that TRPV1^+^ neurons were switched from silent to active states in response to FUS stimulation, while nearby neurons without TRPV1 overexpression were not affected (Fig.3c). We consistently observed FUS activation of the TRPV1^+^ neurons, showing a strong Ca^2+^ influx (Fig. 3e, right panel, and Fig. S5) and a high success rate of 80.8 ± 9.0% (Fig. 3f). For comparison purposes, we randomly selected 13 neurons from mice in group two with the overexpression of mCherry without TRPV1 (TRPV1^-^). In these mice, FUS failed to activate these selected neurons as seen from the minimal Ca^2+^ influx (Fig. 3e, left panel) and a low success rate of 10.7 ± 5.7% (Fig. 3f), which was not significantly different from the control without FUS stimulation (Fig. 3d, 3f). These findings demonstrated the ability of sonogenetics to manipulate the activity of a selected type of neurons in the mammalian brain via TRPV1.

One practical consideration with thermal-based neuromodulation tools is the risk of damaging effects from the temperature increase. Two additional groups of normal mice without the injection of the viral vectors were used to evaluate the safety of FUS exposure (n=4 for each group). One group was sacrificed after FUS sonication with identical parameters as those used in the above experiment (FUS^+^). The other group was used as the control without FUS exposure (FUS^-^). We inspected neuronal integrity, inflammation, and apoptosis using immunohistochemical staining of the neurons (NeuN), astrocytes (GFAP), microglia (Iba1), and cell death (Caspase-3 and TUNEL). Cell nuclei appeared to be intact and the morphology of neurons appeared normal. We did not find a significant change in the numbers of neurons, astrocytes, microglia, and apoptotic cells when comparing FUS^+^ and FUS^-^ groups.

### Precise control of neural activity

Precise control of neural activity via sonogenetics depends on well-defined temporal and spatial control of the ultrasound energy. All the above experiments were performed using a 15 s-long FUS pulsed wave. After demonstrating that the 15 s-long FUS pulsed wave can robustly activate TRPV1^+^ neurons (Fig. 3), we compared pulsed wave (PW) with continuous wave (CW) for targeted activation of TRPV1^+^ neurons. The experimental method was identical to the above *in vivo* study with one modification. Considering the relatively low co-expression rate of TRPV1 and GCaMP6f using the transgenic mice, we switched to co-injection of two viruses: one for the delivery of GCaMP6f and one for the delivery of TRPV1.

Short on-and off-time delays of neuromodulation permit multiple simulations in short observation times, which is crucial for precise correlation of neural activity with brain function and for obtaining statistically significant data. We tested whether we could shorten the stimulation duration by using PW FUS with 7 s total sonication duration and CW FUS with 7, 4, and 1 s sonication duration. Each condition was tested on 5 TRPV1^+^ neurons and each neuron was repeatedly stimulated 10 times. We found that reducing the PW duration to 7 s failed to evoke Ca^2+^ influx robustly, but CW with 7 s and 4 s duration repeatedly activated TRPV1^+^ neurons in the mouse brain *in vivo* (Fig. 4a, Supplementary Video 1). Further decreasing the CW duration to 1 s did not consistently induce activation of the neurons. The success rate of 7-s CW and 4-s CW (88.0 ± 7.3% and 68.0 ± 11.6% respectively) were comparable to 15-s PW stimulation (80.8 ± 9.0%) with no significant difference (Fig. 4b). The onset latencies were 2.6 ± 0.2 s and 2.1 ± 0.3 s for the 7-s CW and 4-s CW stimulation, respectively (Fig. 4c). They were significantly faster than the latency for the 15-s PW (5.9 ± 1.0 s). In addition to faster onset time, CW also reduced the 50% relaxation time. The normalized fluorescence intensity (ΔF/F) was back to half of its peak intensity within 14.6 ± 1.0 s and 15.3 ± 2.0 s for the 7 s and 4 s stimulation, respectively (Fig. 4d).

**Fig. 4.**
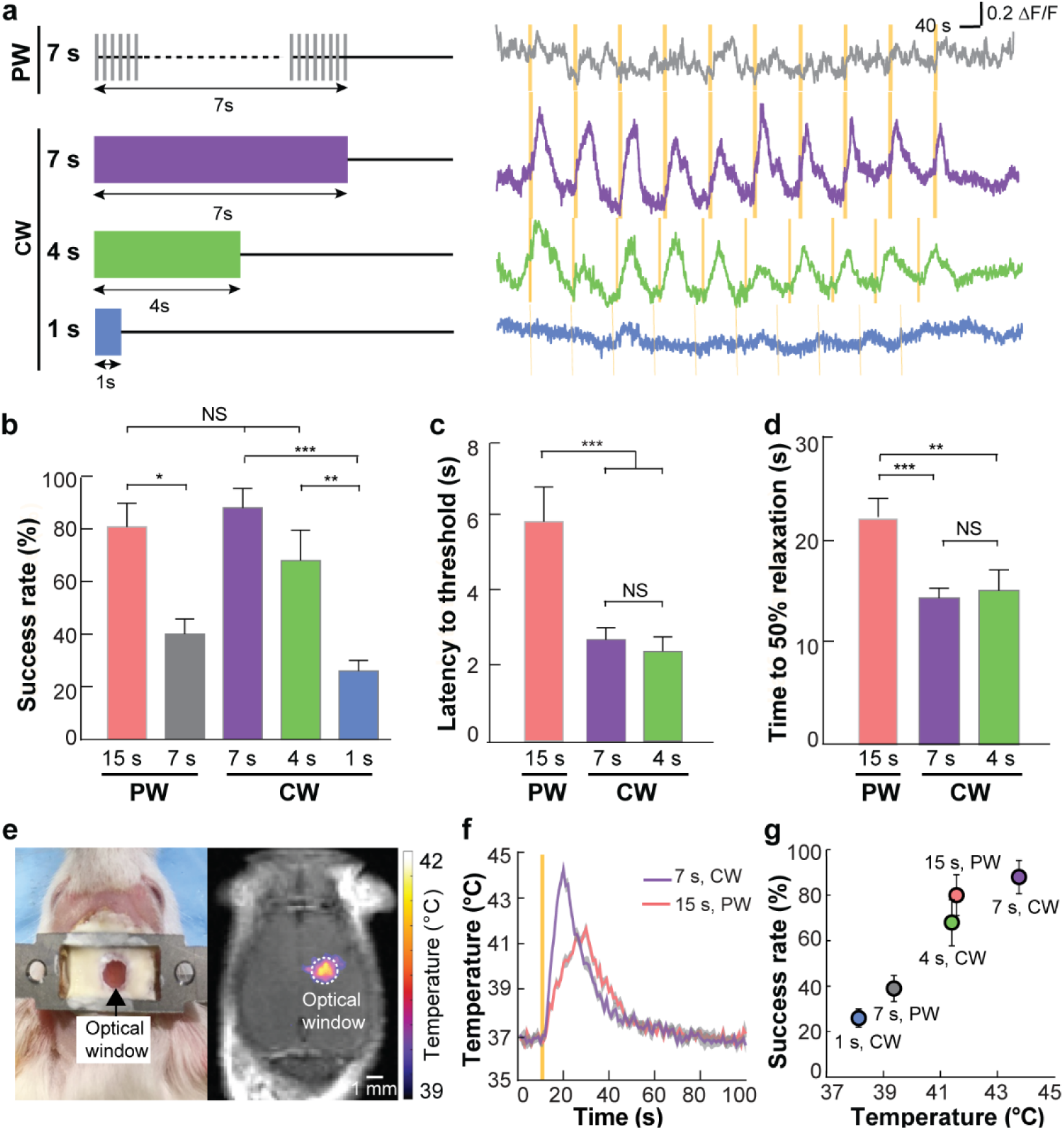
Precise control of neural activity by sonogenetics. (**a**) Illustration of different FUS parameters for stimulation (Left). The representative Ca^2+^ dynamics in response to 10 repeated FUS stimulations using different parameters (Right). Yellow bars indicate the FUS stimulation time. PW: pulsed wave. CW: continuous wave. Quantification of (**b**) Success rate, (**c**) latency, and (**d**) time to 50% relaxation for different FUS parameters (error bars indicate SEM). *: *P*<0.05; **: *P*<0.002; ***: *P*<0.001. (**e**) Spatial distribution of FUS-induced heating as imaged by MRI thermometry *in vivo.* The dashed circle indicates the location of the optical window. An image of the mouse with the optical window is shown on the left. (**f**) Mean temperature within the optical window as a function of time for both PW and CW stimulation. The yellow solid line indicates the onset of FUS stimulation. (**g**) Correlation between the peak temperature associated with each FUS parameter and the success rate of TRPV1^+^ neuron activation. Error bars indicate SEM.

We performed *in vivo* real-time temperature imaging using magnetic resonance (MR) thermometry. MR thermometry is an established technique for noninvasive, real-time, volumetric, and quantitative temperature measurements. The most reliable and commonly used MR thermometry method is based on temperature-sensitive proton resonance frequency shift. We have established the MR thermometry technique for monitoring heating by FUS in our previous studies^24, 25^. MR thermometry shows localized heating at the FUS spatially targeted brain location. The small FUS beam size (Fig. 3a) allows us to precisely target at the cortex underneath the optical window. We found that heat induced by FUS was spatially localized at the targeted brain location (Fig. 4e). We quantified the average temperature change within the optical window for both PW-FUS and CW-FUS stimulation (Fig. 4f). We then calculated the peak temperature associated with each sonication condition and found that the success rate of TRPV1^+^ neuron activation was positively correlated with the peak temperature (Fig. 4g). This positive correlation between temperature and success rate further supports our hypothesis that FUS-induced heating is the dominant mechanism for TRPV1-mediated sonogenetics. The temperature curves also verified that CW FUS raised the temperature faster than PW FUS (Fig. 4f), which could explain the shortened latencies associated with 7-s CW and 4-s CW compared with 15-s PW. These findings suggest that we can control the success rate, on-off temporal dynamics, and spatial location of sonogenetic neuromodulation by using different FUS parameters to effectively control the heating effect.

## Discussion

This study demonstrated that TRPV1-based sonogenetics is a noninvasive, safe, cell-type specific, and spatiotemporally controllable neuromodulation tool. ThermoTRP channels are cation channels of the Transient Receptor Potential family whose conductances change dramatically with temperature^18^. Among the identified thermoTRPs, TRPV1 is the most suitable one for the mammalian application, because its temperature sensitivity threshold is approximately 42 °C^17^. TRPV1 has been used by magnetothermal genetics to modulate neural activity^19, 20^ and control animal behavior^26, 27^. Alternating magnetic fields were used to heat superparamagnetic nanoparticles adhering to the neuronal membrane for the activation of neurons heat-sensitized by expressing TRPV1. Magnetothermal genetics allows noninvasive delivery of energy to control neural activity remotely, but major safety issues arise from the exogenous superparamagnetic nanoparticles incorporated into the brain and poor spatial resolution of the magnetic field. FUS can remotely warm up the brain tissue under precise spatial and temporal control without the need of injecting nanoparticles to the brain. The strong positive correlation between temperature and the success rate of TRPV1^+^ neuron activation (Fig. 4g) has not been reported before by magnetothermal genetics. Our findings that the success rate and the on-off temporal dynamics of TRPV1-based sonogenetics can be precisely controlled by FUS suggest that our proposed sonogenetics is a controllable and reliable neuromodulation tool. Past research effort has been focused on developing sonogenetics based on mechanosensitive ion channels. One most recent study showed that an engineered auditory-sensing protein, prestin, successfully stimulated neurons in deep regions of mouse brains as evaluated by c-fos staining of ex vivo brain slices^16^. However, FUS-evoked calcium influx in HEK293T cells expressing this protein lasted for more than 100 s after a single FUS stimulation with no evidence to show the calcium influx could be turned off. Such poor temporal response limits the application of this protein as a sonogenetic tool and the lack of understanding of how this protein senses ultrasound further hinders its utilization in neuromodulation. Our study opens new avenues for developing sonogentics using thermosensitive ion channels. Future studies are needed to screen other candidate thermo-and mechano-sensitive ion channels to further expand the toolbox for sonogenetics.

Although ultrasound is well-known to be associated with both mechanical and thermal effects, existing efforts on ultrasound neuromodulation focus on harnessing the mechanical effects of ultrasound while avoiding the heating effects. Heating effects have been intentionally avoided for the major concern of tissue destruction as the gap between normal body temperature and the noxious, tissue-damaging temperature is narrow^18^. However, technological advancement has made it possible to real-time monitor and control FUS-induced heating in the human brain based on MR thermometry^28^. Moreover, MR investigation of FUS-induced brain tissue damage suggested a safety thermal dose threshold of about 1 equivalent min at 43 °C for the brain^29^. The thermal doses associated with our FUS parameters were within the range of 0.11 – 0.35 equivalent min at 43 °C, which were much lower than the above safety threshold. The safety of the FUS stimulation was confirmed by our inspection of neuronal integrity, apoptosis, and inflammation markers. We do not rule out potential damage by other FUS parameters or chronic effects occurring over a longer time. Further studies are needed to fully establish the safe operation parameter space of the proposed sonogenetic technique.

The sonogenetic tool based on TRPV1 has a latency of several seconds. This time scale is not limited by the intrinsic properties of the ion channel, as the mammalian TRPV1 responds with a time constant of ~5 ms to an infrared-laser triggered temperature jump, comparable to ligand-or voltage-gated ion channels^30^. This time scale reflects the kinetics of FUS tissue heating to reach the targeted temperature for neuron activation. The latency is longer than optogenetics which has a millisecond temporal resolution. Millisecond temporal resolution is unnecessary for many applications, for example linking activation of specific neurons to behavior outputs^18^. Moreover, the second-scale latency by the sonogenetics is comparable to the response time of magnetothermal neural stimulation^27^ but much faster than chemical stimulation using designer receptors exclusively activated by designer drugs (DREADDs)^31^.

Sonogenetics, like all other genetic-based neuromodulation techniques, requires gene therapy for genetic targeting of specific cell types to express ultrasound-sensitive ion channels. Gene therapy methods based on viral and other vectors have already provided promising results in clinical trials, and new genome editing technologies have brought safe gene delivery a step closer to clinical practice^32^. Recently, several preclinical studies have shown that FUS in combination with microbubbles can transiently enhance the blood-brain barrier permeability for the noninvasive and localized delivery of intravenously injected AAVs to the mouse brain for optogenetics^33^ and chemogenetics^34^. Currently, clinical trials are evaluating the feasibility and safety of the FUS technique for drug delivery to the brain^35–37^. FUS has the promise to achieve localized and noninvasive delivery of the viral vectors encoding ultrasound-sensitive ion channels and activation of the expressed ion channels for fully noninvasive cell-type-specific neuromodulation.

Sonogenetics can accelerate achieving the ultimate goal of neuroscience in understanding intact neural circuits and enable potential clinical applications. Compared with the existing FUS neuromodulation technique, which uses ultrasound alone to activate or inhibit brain regions, sonogenetics can target a specific type of neurons in the chosen brain area. This specificity allows sonogenetics to be used to study of causal relationships between neural activity and a behavioral outcome in a temporally precise manner. Further, optogenetics often requires surgery for the implantation of optical fibers to the brain, which is one of the major challenges in the engineering of optogenetics systems for clinical use. FUS energy can be noninvasively delivered to targeted brain locations. The design of the FUS transducer can be modified to suit different applications. For example, the geometry of the transducer can be designed to generate a focal zone that closely matches the volume of the targeted brain tissue. Moreover, instead of using a single-element FUS transducer, a phased array transducer with multiple elements can be used to form various focus patterns^38^ for multiple-location simultaneous stimulation. The phased array transducer can also electronically steer the beam for sequential stimulation of multiple brain locations. Devices for transcranial delivery of FUS energy for clinical thermal ablation treatment of essential tremor has been approved by the U. S. Food and Drug Administration^39^. We envision that the sonogenetic technique could be extended to large animals and even humans owing to the deep penetration depth of FUS.

Not only will sonogenetic have a broad range of applications in basic and translational neuroscience, its principle of using FUS for noninvasive, spatiotemporal control of biological systems will also impact other fields in biological science and biomedical engineering. In the future, such a technique may even support new therapeutic modalities for humans, enabling the control of cells not only in the brain but also in other parts of the body.

## Methods

### Plasmid and virus

pLenti-CaMKII-TRPV1-p2A-mCherry-WPRE was a gift from P. Anikeeva (MIT). Lentiviral vector was produced according to the established protocol by Hope Center Core at Washington University in St. Louis. Briefly, HEK293T cells were plated at 30–40% confluence 24 h before transfection (70–80% confluence at the time of transfection). 10 g of the lentiviral vector with the appropriate insert, 5.8 µg of pMDLg/pRRE, 3.1 µg of pCMV-G, and 2.5 µg of pRSV-rev were cotransfected into HEK293T cells using the calcium phosphate precipitation method. The medium was replaced with fresh medium containing 6 mM sodium butyrate at 6 h after transfection. Culture supernatant was collected 42 h after transfection. The supernatant was passed through a 0.45 µm SFCA syringe filter (Corning) and ultracentrifuged through a 5 ml 20% sucrose (in PBS) cushion in a polyallomer tube at 11000 rpm, 4 °C, for 4 h with SW28 rotor (Beckman Coulter). The concentrated vector was stored at −80 °C until use. The vector titers were determined by real-time PCR as previously described^40^. To make pLenti-CaMKII-mCherry-WPRE, the mCherry fragment was ligated to pLenti-CaMKII-TRPV1-p2A-mCherry-WPRE in which TRPV1-p2A-mCherry was removed by digestion with BamHI and EcoRI (blunt end-treated).

### *In vitro* cell culture experiments

The HEK293T cell line was a gift from X. W. Wang (Washington University in St. Louis). HEK293T cells were grown in DMEM (4.5 g/*L* glucose), supplemented with 10% FBS, 2 mM L-glutamine, 100 μg/mL sodium pyruvate, 1% non-essential amino acids, and 1% PenStrep. 4×10^5^ HEK293T cells seeded in a 6-well plate 1-day prior were transduced overnight with 2 μL lentivirus of pLenti-CaMKII-mCherry-WPRE or pLenti-CaMKII-TRPV1-p2A-mCherry-WPRE in the presence of 8 μg/ml polybrene. One week later, pLenti-CaMKII-mCherry-WPRE and pLenti-CaMKII-TRPV1-p2A-mCherry-WPRE transduced HEK293T cells were enriched for mCherry positive cells (>95%) using fluorescence-activated cell sorting (FACS) at Siteman Flow Cytometry Core at Washington University in St. Louis. Sorted transfected cells were expanded and used for Ca^2+^ imaging and FUS stimulation.

Fluo-4 AM (Thermo Fisher Scientific), a Ca^2+^ indicator, was used to image the spatial dynamics of Ca^2+^ response to FUS stimulation using a fluorescence microscope (LX70, Olympus). First, a 50 μg Fluo-4 stock solution was diluted by 45 μL Pluronic F-127 20% solution in DMSO (P3000MP, Thermo Fisher Scientific) and added to the HEK293T cells in a 48-well plate to the final concentration of 4 μM. After 45-min incubation at 37 ^°^C, cells were washed in indicator-free live cell imaging solution (Thermo Fisher Scientific) twice and incubated for another 30 min for complete de-esterification. Then the cells were immersed in the live cell imaging solution and the Ca^2+^ images were recorded by the fluorescence microscope at a frame rate of 2.4 frames/sec.

FUS stimulation was applied to the cells during Ca^2+^ imaging using a customized *in vitro* FUS system (Supplementary Fig. S1). The ring-shaped transducer was aligned confocally with the objective of the fluorescence microscope. Above the ring transducer were a layer of 0.2 mm transparent plastic window and a thin layer of degassed ultrasound coupling gel. The wells were filled with the live-cell imaging solution and covered by ultrasound absorber on top (Aptflex F28P, Precision Acoustics) to minimize standing waves. The acoustic pressure output of the FUS transducer was carefully calibrated using hydrophones (HGL-200 and HNP-200, Onda) in a degassed water tank. The FUS parameters applied in the study were: center frequency of 1.7 MHz, peak negative pressure of 1 MPa, duty cycle of 40%, pulse repetition frequency of 10 Hz, and duration of 30 s. The spatial peak temporal average intensity (I_spta_) was calculated to be 12.5 W/cm^2^. The temperature of the imaging solution was monitored using a fiber-optic thermometer (Luxtron, now LumaSense Technologies). The tip of the fiber-optic thermometer was inserted at a location about 1 mm away from the center of the microscope’s FOV for real-time temperature recording. During the experiment, the baseline temperature of the medium was maintained at 37^°^C by a resistor-based heating unit (Caddock File). For the positive control experiment, the cells were heated by water-bath heating using the resistor-based heating unit.

FITC Annexin V (AV) and propidium iodide (PI) were used to assess the effect of FUS stimulation on cell viability. TRPV1^-^ and TRPV1^+^ HEK293T cells were seeded in a 48-well plate and treated by FUS. At 12 h, 24 h, and 48 h after sonication, cells were harvested and stained with AV and PI according to manufacturer’s manual (FITC Annexin V Apoptosis Detection Kit with PI; Biolegend). Stained cells were immediately analyzed by flow cytometry (MACSQuant, Miltenyi Biotec) and the data were analyzed using Flowjo 8.7 software (Tree Star Inc., Ashland) to quantify the percentages of necrotic cells (AV-, PI+), early apoptotic cells (AV+, PI-), late apoptotic cells (AV+, PI+), and live cells (AV-, PI-).

### Intracranial injection

Thy1-GCaMP6f mice (male, 8-week old, Jackson Laboratory) and wild-type BALB/c mice (female, 8-week old, Jackson Laboratory) were subjected to intracranial injection of viral vectors. All animal studies were performed in compliance with guidelines set forth by the NIH Office of Laboratory Animal Welfare and approved by the Washington University Institutional Animal Care and Use Committee. All mouse surgeries were performed under aseptic conditions. Mice were anesthetized via intramuscular injection of ketamine (0.10 mg/g body weight) and xylazine (0.01 mg/g body weight) mixture. Before the virus injection, buprenorphine (buprenex, 0.1 μg/g body weight) and carprofen (Rimadyl, 5 μg/g body weight) were injected subcutaneously. Coordinates used for intracranial injection into the primary somatosensory cortex were determined according to the mouse brain Atlas as follows: −0.5 mm dorsoventral, −1.2 mm anterior-posterior, and −1.2 mm mediolateral. Equivalent dose of lentivirus 1.0 μL of pLenti-CaMKII-TRPV1-p2A-mCherry-WPRE solution or 0.64 μL pLenti-CaMKII-mCherry-WPRE) was injected into the somatosensory cortex of Thy1-GCaMP6f mice using a microinjector (Nanoject II; Drummond Scientific) at a speed of 0.69 μL /min. After the injection, the needle was slowly withdrawn at a speed about 0.5 mm/min. The scalp was closed with vetbond (3M) and sutured, and the mice were allowed to recover on a heating pad. Following the surgery, anti-inflammatory and antibiotic drugs, including sulfamethoxazole (1 mg/ml), trimethoprim (0.2 mg/ml), and carprofen (0.1 mg/ml), were chronically administered in the drinking water. Following the same procedures, BALB/c mice were injected with a mixture of 1.0 μL Lentivirus (pLenti-CaMKII-TRPV1-p2A-mCherry-WPRE) and 0.1 μL of adeno-associated virus serotype 9 (AAV9) carrying GCaMP6f under the excitatory neuronal promoter CaMKII (AAV-CaMKII-GCaMP6f) (Addgene) to co-transfect the TRPV1 and GCaMP6f into neurons.

### *In vivo* 2PM imaging and FUS brain stimulation

4–6 weeks following viral injection, mice were used for FUS stimulation with simultaneous *in vivo* 2PM imaging to record the neural activity. Before the experiment, a chronic cranial window was created on the mouse head to obtain optical access to the mouse cerebral cortex for time-lapse Ca^2+^ imaging using 2PM following an established protocol^41^. Briefly, a small piece of the skull (a circular shape with ~2.3 mm diameter) was removed by a trephine drill (Fine Science Tools). Then 1.2% agarose (Sigma) in artificial cerebrospinal fluid (aCSF, Tocris Bioscience) was applied onto the dura and covered by a 3.5 mm cover glass (#1 thickness, CS-3.5R, Warner instruments). The cover glass was then sealed by vetbond and adhesive cement (C&B-Metabond). Finally, a customized titanium frame was embedded on top of the skull and fixed by adhesive cement. After the optical window surgery, the mice were prepared for 2PM imaging and FUS stimulation. The mice were anesthetized via intramuscular injection of ketamine (0.10 mg/g body weight) and xylazine (0.01 mg/g body weight) mixture. Then, the head was fixed by screwing the titanium frame on top of the mouse’s head to a titanium plate on the 2PM stage (Fig. 3b). Body temperature was kept around normal physiological status (37 °C) by an automatic temperature controller (TC-334B, Warner Instruments).

Time-lapse 2PM imaging was performed with a custom-built two-photon microscope^42^ running SlideBook acquisition software (Intelligent Imaging Innovations) and equipped with a Thor resonant scan head, 2 Bialkali and 2 multi-alkali photomultiplier tubes (Hamamatsu), a 1.0 NA 20× water dipping objective (Olympus), and a Pockel’s cell laser attenuator (Conoptics). Fluorescence was excited with a Chameleon Vision II Ti: Sapphire laser (Coherent) tuned to 920 nm for GCaMP6f and 980 nm for mCherry; fluorescence emission was detected by photomultiplier tubes simultaneously using 495 nm and 575 nm dichroic filters: green (495–575 nm, GCaMP6f) and red (>575 nm, mCherry). The output laser power was set to around 60 mW for Thy1-GCaMP6f mice and 20 mW for AAV-CaMKII-GCaMP6f transfected mice. The image resolution was 0.585 µm/pixel. The original scanning frame rate was 30 Hz and every 10 frames were averaged to generate time-lapse images with 3 Hz frame rate.

The FUS transducer was specially designed in the way that the inner edge of the ring FUS transducer geometrically fit the outer edge of the microscope objective to confocally align the optical beam and FUS beam. In the repeated FUS stimulation studies, the interval between every adjacent stimulation was 80 s to minimize interference among repeated stimulations. A total of 5 different parameter groups were used in the *in vivo* study with the ultrasound frequency (1.7 MHz) and peak negative pressure (1.3 MPa) kept the same and changing the duty cycle and sonication duration: (1) PW with duty cycle of 40% and duration of 15 s; (2) PW with duty cycle of 40% and duration of 7 s; (3) CW with duty cycle of 100% and duration of 7 s; (4) CW with duty cycle of 100% and duration of 4 s; (5) CW with duty cycle of 100% and duration of 1 s;

### Calcium data analysis

The recorded Ca^2+^ images from both *in vitro* cell culture and *in vivo* mice studies were analyzed by Matlab using a published algorithm^43^. For the *in vitro* experiment, cells were automatically identified after applying a constrained nonnegative matrix factorization (CNMF) framework. Then, 100 cells are randomly selected from 3 independent trials. Relative fluorescence intensity changes were computed for calcium signal as ΔF/F=(F-F_0_)/F_0_, where F_0_ represents the average of a 1.5 s-long fluorescent signal acquired before FUS onset. Successful FUS stimulation was defined by the criteria that the normalized Ca^2+^ fluorescence intensity (ΔF/F) acquired from the onset of FUS stimulation to 1.5 s after FUS was both >0.1 and > 2× standard deviation (SD) of 1.5 s-long signal acquired before FUS^44^. The percentage of responsive cells was calculated by dividing successfully stimulated cells over all the selected cells.

For the *in vivo* study, ROIs were manually selected to cover individual soma of the neurons expressing both GCaMP6f and mCherry with TRPV1 for TRPV1^+^ neurons and without TRPV1 for TRPV1^-^ neurons. Successful FUS stimulation was defined as the above *in vitro* study (ΔF/F > 0.1 and >2 × SD). The success rate was quantified by the proportion of successful FUS stimulation to all the stimulation in every single neuron. The statistical result was then calculated by averaging the success rate over all mCherry and GCaMP6f double-positive neurons. Latency to threshold was defined as the time from the onset of FUS to the time of successful stimulation, which was defined above. Time to 50% relaxation was defined as the time from the time reaching the peak amplitude to the time decaying to half of the peak amplitude.

### *In vivo* temperature imaging

MR thermometry was used for non-invasively imaging the spatiotemporal distribution of FUS-induced temperature rise in mice brain *in vivo.* BALB/c mice without viral injection were used in this study. MR thermometry was performed using a (Agilent/Varian DirectDrive Console) 4.7 T MRI system. Temperature images were acquired using a continuously applied gradient-echo imaging sequence with a flip angle of 20 degrees, *T*_R_ of 10 ms and *T*_E_ of 4 ms, slice thickness of 1.5 mm, and matrix size of 128 × 128 for 60 × 60 mm FOV. Phase images were processed in real-time using ThermoGuide software (Image Guided Therapy). MRI compatible FUS transducer (Image Guided Therapy) was targeted at the same brain location as 2PM study and used the same FUS parameters as well, except a slight difference in the ultrasound frequency (1.5 MHz instead of 1.7 MHz), which was restricted by the available transducer. Before the *in vivo* experiment, the performance of the MR thermometry was calibrated in a gelatin phantom by comparing it to the temperature measured by the fiber-optic thermometer used as a gold standard. The difference in the result of these two methods was within 10%.

### Histological analysis

Safety of FUS stimulation was assessed by evaluating the number of neurons, astrocytes, microglia and apoptotic cells using immunohistochemistry analysis. First, 5 sequential FUS stimuli were applied to the BALB/c mice without viral injection using the same FUS system as the above MR thermometry study and the same FUS parameters. At around 2 h after FUS sonication, mice were sacrificed by transcardial perfusion with 4% paraformaldehyde (PFA). Brains were harvested and fixed in 4% PFA overnight and equilibrated in 30% sucrose for cryosectioning. The fixed brains were sectioned to 20 µm slices. Then, the slices were preprocessed in 0.3% v/v Triton X-100 and 3% v/v blocking serum solution in PBS for 1 h in the dark at room temperature to increase the permeability and block the background. Then the slices were washed 3 times using PBS and incubated in primary antibody solution with 3% blocking serum overnight at 4 °C. The primary antibodies used in this study include: anti-NeuN (Abcam, Cat: 104225, 1:1000), anti-GFAP (Abcam, Cat: 07165, 1:1000), anti-Iba1 (Abcam, Cat: 178846, 1:1000,), anti-Caspase-3 (Cell Signal, Cat: 9661s, 1:1000), and DeadEnd™ colorimetric TUNEL System (Promega, Cat: G7360). After 3 PBS wash, the slices were then incubated in secondary antibody (Donkey antirabbit Alexa Fluor 488, Jackson laboratory, 1:400) in the dark for 3 h at room temperature. Finally, the slices were moved onto glass slides and mounted with VECTASHIELD (Vector Laboratories) that contained DAPI nuclear stain.

### Statistical analysis

Data were analyzed using either a two-tailed t-test with unequal variance (in Fig. 3h) or two-way ANOVA with Bonferroni post hoc test (when more than two samples were compared in Fig 2d, 3f). One-way ANOVA with Bonferroni post hoc test was used in Fig 4b, 4c, and 4d. All data with *p* < 0.05 were considered to be significant.

### Data availability

The data that support the findings of this study are available from the corresponding authors upon reasonable request.

## Acknowledgments

This work was supported by the National Institutes of Health (NIH) BRAIN Initiative (R01MH116981) and NIBIB (R01EB027223). We thank Dr. Hunter Banks, Dr. Lu Zhoa, and Ms. Lindsey Brier for the insightful discussions. We also thank D. Mingjie Li from the Hope Center Viral Vectors Core at WashU for preparing the lentiviral and AAV vectors. We also appreciate the help of Dr. James D. Quirk in thermometry mapping using magnetic resonance imaging.

## Author contributions

H.C. and Y. Y. designed the experiments. Y. Y. conducted the experiment with the assistant of other co-authors. J.Y. assisted with the *in vitro* experiment. D.Y and M.M assisted with setting up the two-photon microscopy system. L.Z and C.P assisted with the MR thermometry experiment. H.B and Y.Y assisted with animal surgery. Y.Y performed the immunohistochemistry staining experiment. J.C, J. P.C, and M. R.B contributed to the discussion and interpretation of the results. H.C and Y.Y wrote the manuscript and all authors modified and revised the manuscript.

