## Supplementary for "Sonogenetics for noninvasive and cellular-level neuromodulation in rodent brain"

### Supplementary Information

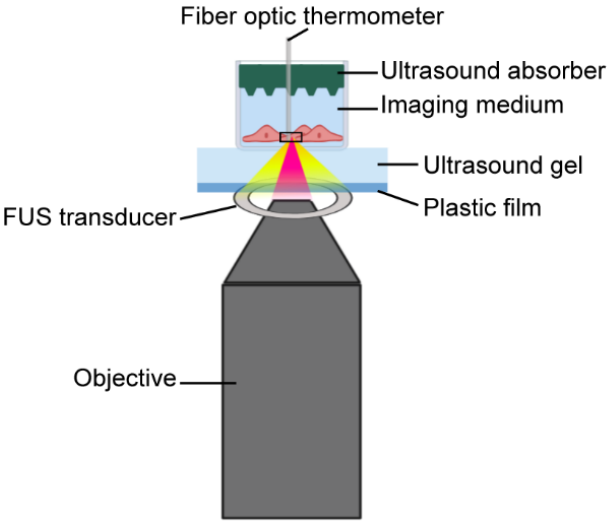


**Fig. S1. Illustration of the experimental setup for *in vitro* simultaneous FUS stimulation and $Ca^{2+}$ fluorescence imaging.**

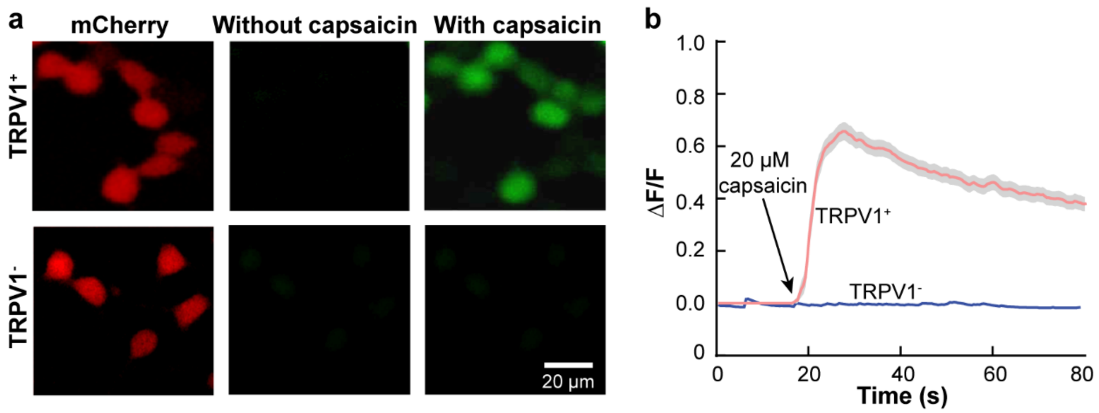


**Fig. S2. Functional assessment of TRPV1 ion channel expressed in HEK293T cells. (a)** Fluorescence images of HEK293T cells transduced with the pLenti-CamKII-TRPV1-p2A-mCherry ($TRPV^+$) or pLenti-CaMKII-mCherry-WPRE ($TRPV^-$). Capsaicin, a TRPV1 agonist, was added to test the functionality of TRPV1. **(b)** $Ca^{2+}$ fluorescence intensity change (ΔF/F) as a function of time. Shaded areas represent standard error of the mean and solid lines are the average intensity of 30 randomly selected cells.

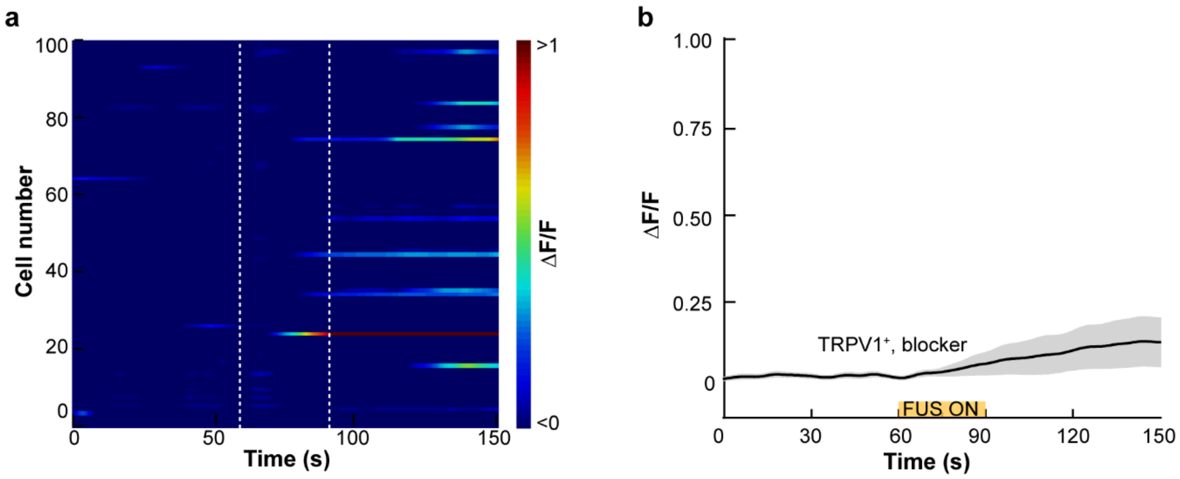


**Fig. S3. TRPV1 antagonist capsazepine reduces FUS-induced $Ca^{2+}$ influx in TRPV1$^+$ HEK293T cell**. (**a**) Heat map of the $Ca^{2+}$ fluorescence intensity of 100 automatically selected TRPV1$^+$ cells treated by adding 20 μM capsazepine (TRPV1 antagonist) followed by FUS stimulation. (**b**) Fluorescence intensity as a function of time of the cells in (a) that responded to FUS stimulation. Shaded area indicates SEM and the solid line indicates the mean intensity.

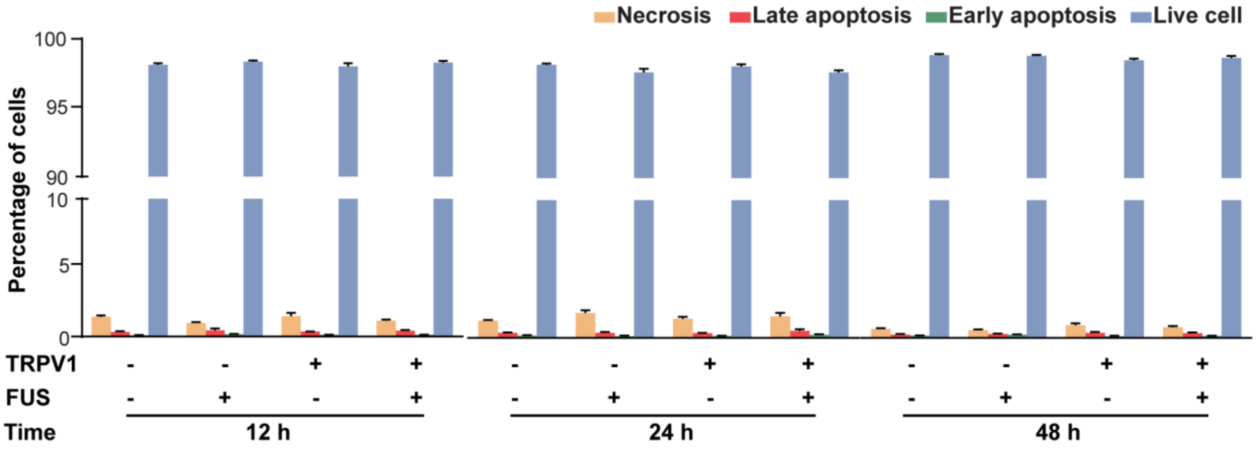


**Fig. S4**. **Evaluation of HEK293T cell viability after FUS stimulation**. The cells were labeled with Annexin V-FITC (AV) and propidium iodide (PI), and then measured by flow cytometry. Percentages of necrotic cells (AV-, PI+), early apoptotic cells (AV+, PI-), late apoptotic cells (AV+, PI+), and live cells (AV-, PI-) were measured at 12 h, 24 h and 48 h after FUS treatment. Error bars indicate SEM for 4 independent trials.

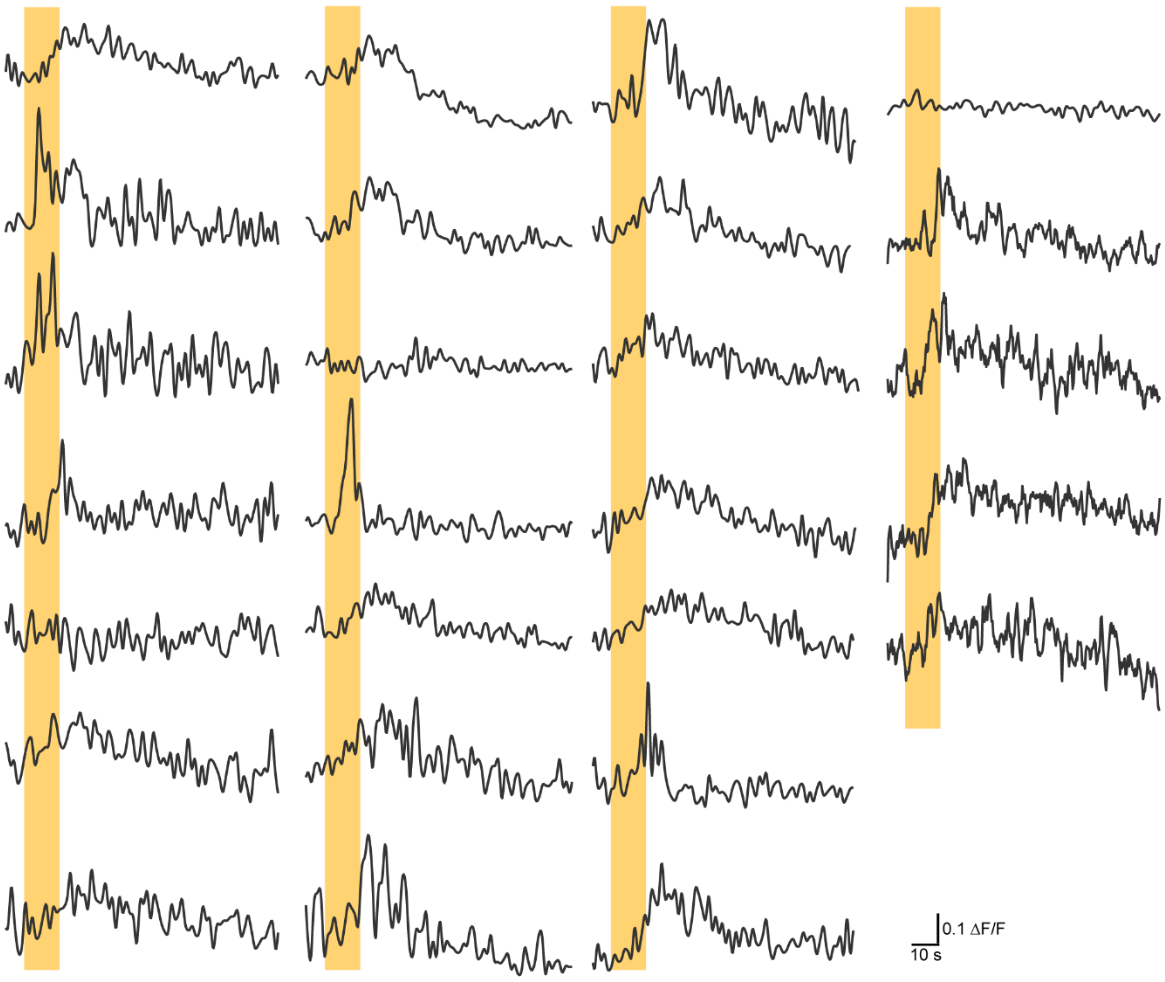


**Fig. S5**. **$Ca^{2+}$ fluorescence intensity curves quantified based on in vivo two-photon microscopic imaging of 26 independent trials from 13 neurons in the mouse cortex**. Yellow bars indicate the FUS stimulation time.
